# Single-cell fluorescence imaging reveals heterogeneity in senescence biomarkers and identifies rapamycin-responsive sub-populations

**DOI:** 10.1101/2024.11.21.624614

**Authors:** Vijayraghavan Seshadri, Charmaine Chng, Joel Tyler, Kaveh Baghaei, Yan Wang, Nuri Gueven, Iman Azimi

## Abstract

Cellular senescence is a state of irreversible cell cycle arrest accompanied by a distinctive inflammatory secretory profile known as the senescence-associated secretory phenotype (SASP). While various biomarkers, such as senescence-associated beta-galactosidase (SA-βgal), EdU incorporation, P21, and P16, are used to identify senescent cells, no single biomarker universally defines cellular senescence, and current methods often fail to address heterogeneity in biomarker expression levels. This study leverages single-cell fluorescence imaging to assess multiple senescence markers including SA-βgal enzymatic activity, P21 and IL-6 expression, and nuclear and cell area, in chemotherapy-induced (mitomycin C) and oxidative stress-induced (D-galactose) senescence models in human fibroblasts.

Our findings reveal significant heterogeneity in SA-βgal activity and distinct sub-populations within senescent cells. Nuclear and cell area measurements emerged as robust indicators of cellular senescence, displaying similar variability across individual cells. Importantly, we identified specific nuclear area sub-populations that strongly correlate with IL-6 expression levels, demonstrating a relationship between the heterogeneous expression of senescence biomarkers and the SASP. To address this heterogeneity, we introduced an induction threshold method to more accurately quantify the percentage of cells expressing senescence biomarkers.

Furthermore, in both senescence models, we observed that rapamycin, a well-known senomorphic agent, selectively targets specific biomarker-expressing sub-populations. This study underscores the value of assessing cellular heterogeneity in senescence research and provides an improved approach for analysing senescence markers in diverse cellular contexts.

## Introduction

Cellular senescence is closely associated with organismal aging and becomes more prevalent with age, especially in organs affected by age-related diseases (1, 2). Senescence involves irreversible growth arrest due to natural ageing or in response to different forms of stress. It is characterised by a heightened secretory phenotype and resistance to apoptosis (3-5). Senescence impacts many different biological processes, including tissue homeostasis, embryonic development, wound healing, immune responses, cell clearance, and cancer (4, 5).

Proliferative senescence was first reported by Hayflick and Moorhead in 1961, who observed that normal cultured human fibroblasts have a finite capacity for cell division under optimal *in vitro* conditions, eventually entering a stage of irreversible growth arrest (1, 3). In addition to natural ageing, cellular senescence can be accelerated by different stressors, such as oxidative stress, oncogene activation, and acute DNA damage caused by chemotherapy or radiotherapy (6, 7). These stressors induce DNA damage that cells cannot fully repair, activating the DNA damage response signalling pathway and leading to senescence (6-9).

Despite their non-proliferative status, senescent cells remain metabolically active and adopt a pro-inflammatory state, releasing cytokines, chemokines, growth factors, and proteases. This collective pro-inflammatory state is termed the senescence-associated secretory phenotype (SASP) (10, 11). Through the SASP, senescent cells communicate with the immune system to either facilitate their clearance or support tissue regeneration. However, this process can also induce senescence in neighbouring healthy cells, contributing to tissue degeneration.

The effects of senescent cells within tissues can be both positive and negative, depending on whether these cells are transient or persistent. Acute senescence plays a beneficial role in tumour suppression, wound healing, and tissue homeostasis, as the SASP signals the immune system to clear senescent cells and promotes the repair of damaged tissue (2, 12). Conversely, with age, the accumulation of senescent cells increases due to the declining efficiency of the aging immune system to remove them (11, 12). This accumulation disrupts tissue structure and function and leads to further induction of senescence. Over time, this build-up, referred to as chronic senescence, results in the persistent release of SASP factors into the microenvironment (11, 12). Consequently, the chronic inflammatory environment induced by these persistent senescent cells becomes harmful, contributing to the progression of various age-related diseases, such as osteoarthritis, atherosclerosis, neurodegenerative disorders, and tumorigenesis (2, 13). Thus, the paradoxical nature of senescence provides short-term benefits but also poses long-term liabilities as senescent cells accumulate over time.

Senescent cells exhibit distinct characteristics that differentiate them from non-senescent cells, providing opportunities for targeted therapeutic interventions. These cells are characterised by specific marker proteins, including the increased expression of cyclin-dependent kinase inhibitors p16, P21, and TP53, which are induced to halt cell proliferation (8, 9, 14). Additionally, senescent cells display unique morphological features, such as enlarged and flattened cellular shapes, irregular and enlarged nuclei, and heightened enzymatic activity of the lysosomal enzyme senescence-associated beta-galactosidase (SA-β-gal), among others (9).

While these features are commonly associated with senescence, they are not exclusive indicators. For instance, the cyclin-dependent kinase inhibitor P21 is not universally upregulated in all senescent cells, making it an unreliable marker when used alone (15). Similarly, p53, known for its role in cellular apoptosis, does not definitively distinguish between senescent and apoptotic cells (16). Furthermore, SA-β-gal, a widely used senescence marker, is also expressed by other cell types, such as macrophages (17, 18). Conditions like high cell density or exposure to hydrogen peroxide can also trigger SA-β-gal activity, leading to false positives in senescence detection (19).

The diversity of senescent cell traits and the lack of a single universal marker necessitate a comprehensive approach that combines multiple senescence biomarkers for precise and reliable identification and quantification of senescent cells. However, this integration presents detection challenges. Traditional methods, such as Western blotting, immunofluorescence, and flow cytometry, often lack the specificity and sensitivity required. While Western blotting can detect specific protein levels, it falls short in providing information on cellular morphology or the spatial distribution of senescence markers and does not capture single-cell heterogeneity in biomarker expression. Immunofluorescence and flow cytometry offer more detailed spatial and quantitative data but struggle due to the nonspecific nature of many senescence markers and the heterogeneity among senescent cell populations. Furthermore, these methods typically involve fixed cells, limiting the ability to study live cell dynamics. Quantitative real-time PCR (qPCR) can be used to quantify mRNA levels of senescence biomarkers; however, it faces challenges in correlating mRNA levels with protein expression. This discrepancy is particularly evident with P21 and p16, where mRNA levels often fail to reach significant thresholds needed for reliable identification of senescence.

Given the current limitations in detecting and analysing senescent cells, there is an urgent need for more robust methods. Existing senescence biomarkers lack specificity and reliability, and no consensus has been reached on their optimal detection. This study aimed to address these limitations by developing a robust method for analysing senescence markers using live single-cell data and high-content fluorescence microscopy. We assessed multiple senescence markers in both live and fixed cells across two distinct models of cellular senescence in human dermal fibroblasts.

## Methods

### Cell culture

Primary human skin fibroblasts (HDFs) (106-05N, Sigma-Aldrich, MO, USA, for Figures 4 and 5; or PCS-201-010, ATCC, for the remaining figures) were cultured in Dulbecco’s Modified Eagle Medium (DMEM, D5523, Sigma-Aldrich) supplemented with 10% foetal bovine serum (FBS). Cells were maintained in a humidified incubator at 37 °C with 5% CO_2_. Cells used for experiments were at passages 5 to 8 (ATCC) and 15 to 19 (Sigma-Aldrich).

For chemotherapy-induced senescence, cells were exposed to various concentrations (50 nM– 600 nM) of Mitomycin C (M4287, Sigma-Aldrich) for 48 hours, followed by a media change to non-Mitomycin C media for an additional five days. For oxidative stress-induced senescence, cells were treated with different concentrations (3.125 mM–200 mM) of D-galactose (D0750, Sigma-Aldrich) for seven days.

For rapamycin treatment, HDFs were pre-incubated with 500 nM rapamycin for 24 hours before the media was replaced with media containing 200 nM MMC + 500 nM rapamycin and incubated for an additional four days. Subsequently, the media was changed to media containing 500 nM rapamycin for a further three days, after which senescence biomarker assays were performed.

### Live single-cell assessment of SA-β-gal

Cells were seeded at a density of 1.5 × 10^6^ in T-25 flasks (430629, Corning). After 48 hours, cells were treated with different concentrations of MMC (50, 100, 200, 400, and 600 nM) 4 for days, after which the media was changed to non-MMC-containing media. For D-galactose-induced senescence, cells were plated at a density of 500 cells per well in glucose-containing media for 24 hours, followed by a media change to glucose-free media with the addition of D-galactose at concentrations of 3.125, 6.25, 12.5, 25, 50, 100, and 200 mM. Seven days post-induction, cells were seeded at a density of 500 cells per well in black 96-well plates (655090, µClear®, Greiner, Germany). After 24 hours, SA-β-gal activity was assessed using the FastCellular™ Senescence SPiDER-βGal Detection Kit (Product Number 092690301, MP Biomedicals, CA, USA).

Following the manufacturer’s protocol, cells were washed with Hank’s Balanced Salt Solution (HBSS) and incubated with Bafilomycin A1 solution for one hour, followed by a 45-minute incubation with SPiDER-βGal, Bafilomycin A1, and 5 µM Hoechst solution (3342, Thermo Fisher Scientific) to stain the nucleus. Cells were then washed twice with HBSS, and plates were prepared for imaging. Images were recorded using high-content cell imaging systems, either the IN Cell Analyzer 2200 (GE Healthcare Life Sciences, IL, USA) at 10x magnification or the Opera Phenix™ Plus (Revvity, MA, USA) at 20x magnification, with excitation at 475 nm and emission at 594 nm. Images were analysed using IN Carta Image Analysis Software (GE Healthcare Life Sciences) or Harmony™ software, version 5.1 (Revvity).

Single-cell mean fluorescence values and total fluorescence (Mean intensity * area of the mask) were quantified by the software. Sub-population data analysis was conducted by binning fluorescence values at intervals specified in the figures, using GraphPad Prism version 9 and above. The threshold induction method was applied by setting a threshold value where at least 90% of values in vehicle-treated cells (0.1% DMSO) fell, and the parameter was reported as the percentage of cells exceeding this threshold. The induction threshold was set at 90% of the values observed in vehicle-treated cells, as 90% of these cells incorporated EdU, while the remaining 10% entered either a quiescent or senescent state. This induction threshold was also applied to the other biomarkers mentioned below.

### Immunofluorescence analysis

To assess P21 ^WAF1/CIP1^ expression, cells were seeded at a density of 500 cells per well in black 96-well plates (655090, µClear®, Greiner). Post seeding (48 h), cells were treated with 200 nM MMC. Seven days post-senescence induction by MMC, cells were fixed with 4% formaldehyde for 10 minutes and permeabilized with 0.5% Triton® X-100 in phosphate-buffered saline (PBS) for 10 minutes. Cells were then blocked with 10% normal goat serum (G9023; Sigma-Aldrich) in PBST (0.1% Tween-20 in PBS) for 1 hour. After blocking, cells were incubated with primary antibodies (P21 WAF1/CIP1 Polyclonal Antibody, Catalog #14-6715-81, eBioscience™, Invitrogen) overnight at 4 °C. Secondary antibodies for P21 (2 µg/ml concentration, Alexa Fluor 647, Catalog #A-21237, Thermo Fisher Scientific) were added to the cells for 1 hour at room temperature (RT). Nuclear staining was performed using Hoechst dye (5 µM). Images were recorded using the Opera Phenix™ Plus (Revvity) at 20x magnification. Harmony™ 5.1 software was used to analyse the images.

### EDU Incorporation

To assess EdU (5-ethynyl 2’-deoxyuridine) incorporation following MMC treatment, cells were treated with 50 and 200 nM MMC. After 48 hours, the cells were cultured in non-MMC-containing media, and on day 5 post-MMC treatment, the media was changed to EdU-containing media (5 µM) for 48 hours. After this incubation, cells were fixed with 4% paraformaldehyde (PFA) for 10 minutes and permeabilized with 0.1% Triton-X100. EdU incorporation was then assessed using the Click-iT™ Plus EdU Cell Proliferation Kit for Imaging with Alexa Fluor™ 488 dye (C10637, Thermo Fisher Scientific).

### Western Blotting

A total of 10 µg of protein was combined with 5 µL of Laemmli buffer (1:9 β-mercaptoethanol [PCS 1610710, Bio-Rad] to Laemmli sample buffer [PCS 161-0737, Bio-Rad]) and sufficient water to reach a final volume of 20 µL. Samples were heated at 95 °C for 5 minutes, then loaded onto either 12% TGX Fast Cast acrylamide gels (PCS 1610174, Bio-Rad) or 4–15% Mini-PROTEAN TGX pre-cast gels (PCS 4561084, Bio-Rad), and subsequently transferred onto PVDF membranes. Membranes were blocked for 1 hour with 5% milk powder in Tris-buffered saline with 0.1% Tween-20 (TBST) and then washed before overnight incubation at 4 °C with primary antibody diluted in 5% milk powder in TBST. The primary antibody used was rabbit anti-P21 (PCS 14-6715-81, Thermo Fisher, MA, USA). After five washes with 5% TBST (5 minutes each), membranes were incubated for 1 hour with secondary antibody goat anti-rabbit IgG (PCS 1706515, Bio-Rad) at a 1:3000 dilution in 5% milk powder in TBST. After washing the membranes five additional times with 5% TBST (5 minutes each), they were incubated in Clarity Western ECL Blotting Substrate (PCS 1705060, Bio-Rad) for 4 minutes. Blot images were captured using the ChemiDoc XRS+ imaging system (Bio-Rad), and densitometry was performed using Image Lab software. Following P21 imaging, membranes were washed five times with 5% TBST (5 minutes each) and incubated for 1 hour at room temperature with mouse anti-β-actin primary antibody (PCS A5441, Sigma-Aldrich, MA, USA) diluted 1:10000 in 5% milk powder in TBST. The membranes were then washed five times in 5% TBST (5 minutes each) and incubated with goat anti-mouse IgG secondary antibody (PCS 1706516, Bio-Rad) at a 1:3000 dilution in 5% milk powder in TBST for 1 hour at room temperature. Following a final set of five washes in 5% TBST (5 minutes each), the membranes were incubated in Clarity Western ECL Blotting Substrate (PCS 1705060, Bio-Rad) for 4 minutes. Blot images were captured using the ChemiDoc XRS+ imaging system (Bio-Rad), and densitometric analysis was performed using Image Lab software.

### Cell Proliferation assay

HDFs treated with MMC and control cells treated with vehicle (0.1% DMSO) were plated at a density of 10^4^ cells per well in black 96-well plates (655090, µClear®, Greiner). Cell proliferation was assessed by measuring fluorescence using the PrestoBlue™ Cell Viability Reagent (Catalog #A13261, Thermo Fisher Scientific) on days 6, 12, and 18, according to the manufacturer’s protocol.

### Statistical analysis

Statistical analyses were performed using GraphPad Prism (version 9.1 and higher, GraphPad Software, CA, USA). Data are reported as mean ± standard deviation (SD), with p values of < 0.05 considered statistically significant. Specific statistical tests and significance values for each experiment are provided in the corresponding figure legends.

## Results

We first aimed to develop a chemotherapy-induced accelerated model of senescence in HDFs using a live-cell fluorescent assay. Our goal was to establish a robust method for senotherapeutic drug discovery based on a universally accepted hallmark of senescence, SA-βgal. To achieve this, we assessed the enzymatic activity and fluorescence intensity of SA-βgal, a key senescence marker, seven days after exposing cells to various concentrations of mitomycin C (MMC) using high-content fluorescence microscopy.

We observed a concentration-dependent increase in SA-βgal enzymatic activity, with a statistically significant three-fold increase beginning at 200 nM MMC treatment (Fig. 1B). To assess the proliferation state of MMC-treated HDFs, we examined EdU incorporation and found a 16-fold decrease in the number of EdU-positive cells compared to controls (Fig. 1C). However, a small sub-population, approximately 5% of MMC-treated HDFs, still incorporated EdU into their nuclei. Given the presence of this small EdU-positive subpopulation, we further investigated whether these cells exhibited enhanced proliferation over time. We conducted a time-course assay to assess proliferation at multiple time points, revealing no increase in proliferation at 12 and 18 days post-MMC treatment, while untreated cells exhibited a two-fold increase in proliferation over the 12 day period (control cells were assessed only until day 12, as they became confluent, leading to contact inhibition) (Fig. 1D).

**Figure 1.**
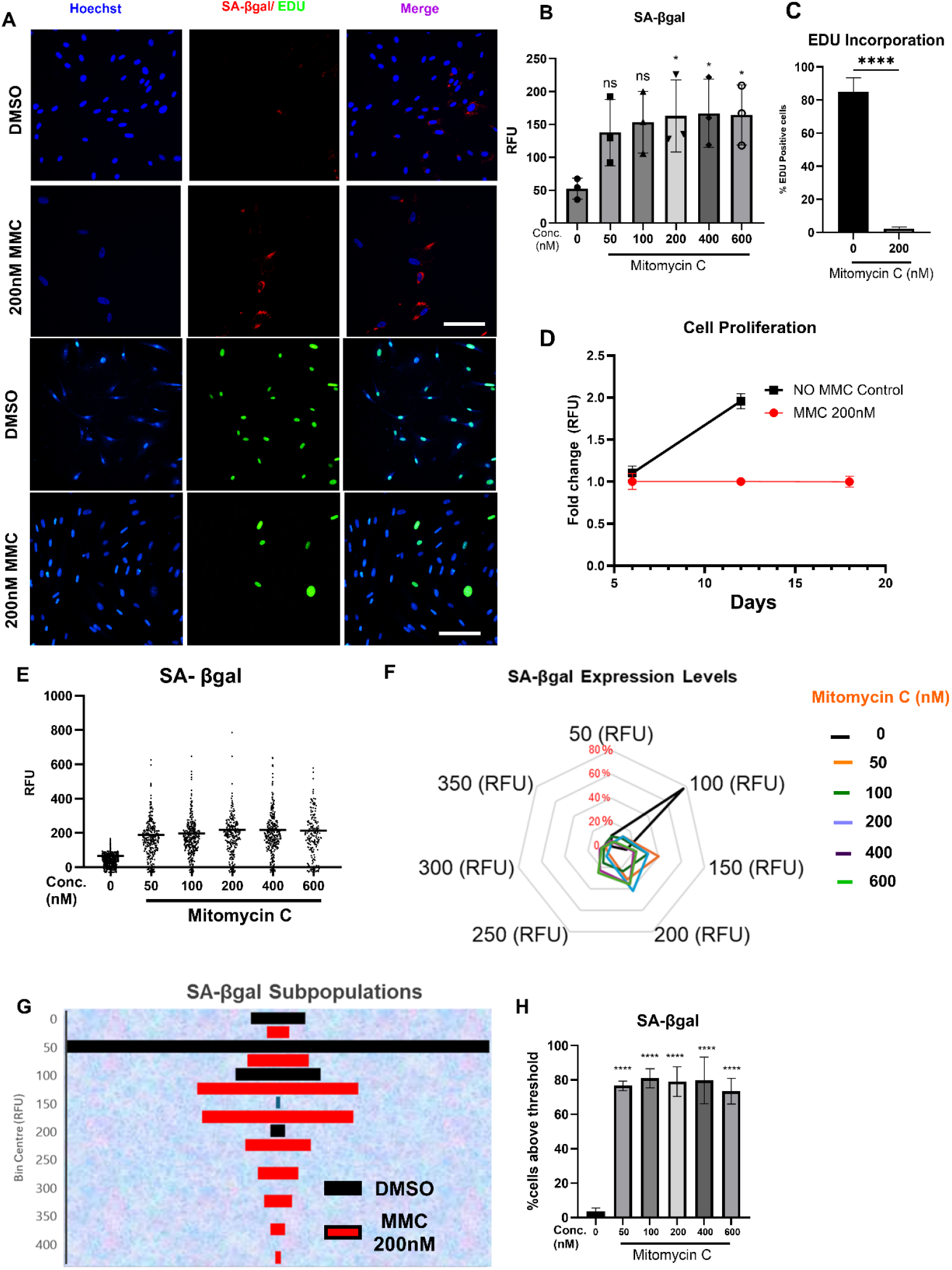
Assessment of SA-βgal and cell proliferation in an MMC-induced senescence model. (**A)** Representative images of SA-βgal staining and EDU incorporation in HDFs treated with vehicle and 200 nM MMC. Images obtained with an IN-Cell analyser 2200 (SA-βgal) and Opera Phenix plus (EDU). Images show nuclei stained with Hoechst (blue), SA-βgal (red)/ EDU (Green) and merged image (scale bar = 100 µm). (**B**) Average fluorescence intensities of SA-βgal in HDFs treated with vehicle (0.1% DMSO) or different concentrations (50 nM-600 nM) of MMC; Error bars represent mean ± standard deviation from three independent biological replicates. * *p* < 0.05 (Simple one-way ANOVA compared with the (0) control group). (**C)** Percentage of cells positive for EDU incorporation in HDFs treated with MMC; \*\*\*\**p* < 0.0001, (Student unpaired t-test compared with 0 group). (**D)** Time course effect of cellular proliferation on HDFs with MMC. **(E)** Single cell fluorescence intensities of SA-βgal in HDFs treated with vehicle (0.1% DMSO) or different concentrations (50 nM-600 nM) of MMC. **(F)** Radar chart of different SA-βgal fluorescence intensities in HDFs treated with vehicle or different concentrations of MMC. **(G)** Sub-population analysis of SA-βgal fluorescence intensities in HDFs treated with vehicle or 200 nM MMC. **(G)** Percentage of cells with SA-βgal intensity of greater than that of the threshold set in the control cells from the respective histograms. Error bars represent mean ± standard deviation from three independent biological replicates. **** *p* < 0.0001 (Simple one-way ANOVA compared with the (0) control group).

Since the SA-βgal assay was performed using a high-content microscope capable of generating single-cell data, we plotted SA-βgal fluorescence intensity levels of individual cells. This analysis revealed significant heterogeneity in SA-βgal activity among MMC-treated cells (Fig. 1E). To further explore this heterogeneity, we performed a sub-population analysis by binning the single-cell fluorescence values of SA-βgal enzymatic activity into specific bin centres (Fig. 1F). We found that the SA-βgal fluorescence values of vehicle-treated HDFs clustered around 100 RFU, while most MMC-treated HDFs (50 nM–600 nM) clustered at 150 and 200 RFU. Further analysis of a single MMC concentration (200 nM) identified distinct sub-populations with unique bin centres for SA-βgal fluorescence intensities (Fig. 1G). This heterogeneity explains why we observed only a three-fold increase in mean fluorescence intensity in MMC-treated groups compared to controls, as shown in Fig. 1B.

To address this heterogeneity, we defined an “induction threshold” for SA-βgal activity based on the fluorescence intensity below which 90% of the control group’s fluorescence values fell (detailed in the Methods section). Cells with fluorescence levels exceeding this threshold were classified as SA-βgal positive. Our analysis showed an approximately 19-fold increase in the percentage of SA-βgal-positive cells in all MMC-treated groups (∼80%) compared to controls (4.2%) (Fig. 1H).

We next extended our investigation to other commonly used senescence biomarkers, including nuclear area, cell area, and P21 expression. MMC treatment resulted in a significant 1.5-to 2-fold increase in the average nuclear area in HDFs (Fig. 2A-B). Similar to the SA-βgal data, considerable variation in nuclear area was observed among individual cells (Fig. 2C). Sub-populations with distinct nuclear areas were evident, as confirmed by differential clustering in the corresponding heatmap (Fig. 2D). Applying our induction threshold method to nuclear area across three independent trials revealed a consistent and robust 6-to 7-fold increase in MMC-treated groups compared to vehicle-treated controls (Fig. 2E).

**Figure 2.**
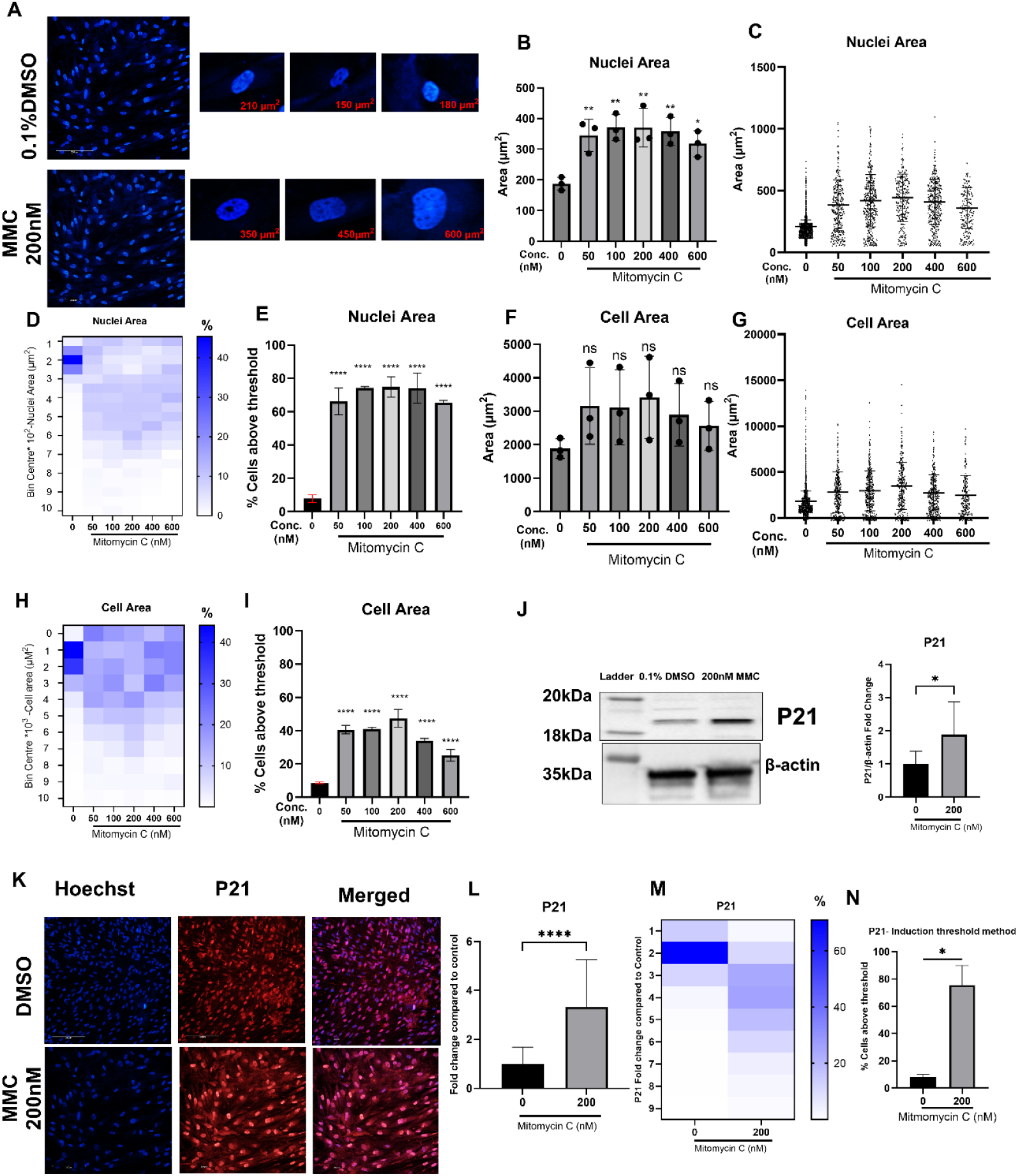
Assessment of nuclear area, cell area and P21 expression in an MMC-induced senescence model in HDFs. Representative images were taken using Opera Phoenix plus ™ at 20X magnification. **(A)** Representative image of Hoechst-stained nuclei in HDFs treated with vehicle and 200 nM MMC treated HDFs. Average value of nuclei area **(B)**, cell area **(F)**, P21 **(L)** in vehicle and MMC treated HDFs; Error bars represent mean ± standard deviation from three independent biological replicates. For Nuclei and cell area (ns, not significant, * *p* < 0.05 ** *p* < 0.01 (one-way ANOVA compared with the (0) control group). For P21 (unpaired t-test compared to the control (0) group **** *p* < 0.0001). Single-cell data for nuclear area **(C)** and cell area **(G)** in HDFs treated with MMC. Individual cell-derived histogram data categorised into various bin centres for nuclear area **(D)**, cell area **(H)** and P21 **(M)** in vehicle- and MMC-treated HDFs. **(J)** Representative image and quantitation of western blot showing the expression of P21 and β-actin, along with the relative quantitiation of expression of P21/β-actin expression in MMC-treated HDFs; \**p* < 0.05, (Student’s unpaired t-test compared with 0 control group). **(K)** Representative images of P21 expression in the nuclei of HDFs treated with vehicle and 200 nM MMC, with nuclei labelled with Hoechst (blue), P21 (red), and merged image (scale bar: 10 µm). Percentage of cells with nuclear area **(E)**, cell area **(I)**, and P21 expression **(N)** exceeding the threshold set in the control cells, derived from the respective heatmaps in vehicle- and MMC-treated HDFs. Error bars represent the mean ± standard deviation from three independent biological replicates. For nuclear area and cell area (**** *p* < 0.0001, one-way ANOVA compared to the control group); For P21 (**** *p* < 0.0001, unpaired t-test compared to the control group).

We also assessed cell area, a cytoskeletal biomarker of senescence. Our results showed a 1.8-fold increase in cell area in HDFs treated with 50-200 nM MMC, though cell area decrease at higher MMC concentrations (Fig. 2F). These increases in cell area were not statistically significant. Nonetheless, single-cell data and heatmap analysis displayed trends similar to those observed for SA-βgal and nuclear area (Fig. 2G). The percent frequency distribution of cell areas revealed the emergence of larger cell populations in MMC-treated groups compared to the control (Fig. 2H). Further analysis using the induction threshold method showed a statistically significant 4.5-fold increase in cell area in cells treated with 200 nM MMC compared to the control group (Fig. 2I).

Another key biomarker for identifying senescent cells both *in vitro* and *in vivo* is P21, a downstream target of TP53 that plays a central role in cell cycle arrest and has various other physiological functions. Initially, we measured P21 expression conventionally using western blot analysis and observed a statistically significant two-fold increase in HDFs treated with 200 nM MMC (Fig. 2J).

Next, we assessed P21 expression at a single-cell level using high-content microscopy (Fig. 2K), which revealed a statistically significant three-fold increase in P21 levels in MMC-treated HDFs. However, further single-cell analysis uncovered substantial heterogeneity in P21 expression, as illustrated by the scattered histogram (Fig. 2M). Applying the induction threshold method showed a 7.5-fold increase in P21 expression in MMC-treated HDFs (Fig. 2N), providing a more robust and accurate assessment of this senescence biomarker compared to the traditional method of averaging values.

To investigate the correlation between different senescent markers and determine if these markers can be used interchangeably, we generated correlation plots between SA-βgal, nuclear area, and cell area (Fig. 3A & B). Positive correlations with r values of around 0.6 for nuclear area (Fig. 3A) and 0.8 for cell area (Fig. 3B) were observed. Notably, these r values were not statistically different between the control and treatment groups, suggesting the presence of heterogenous populations across all groups.

**Figure 3.**
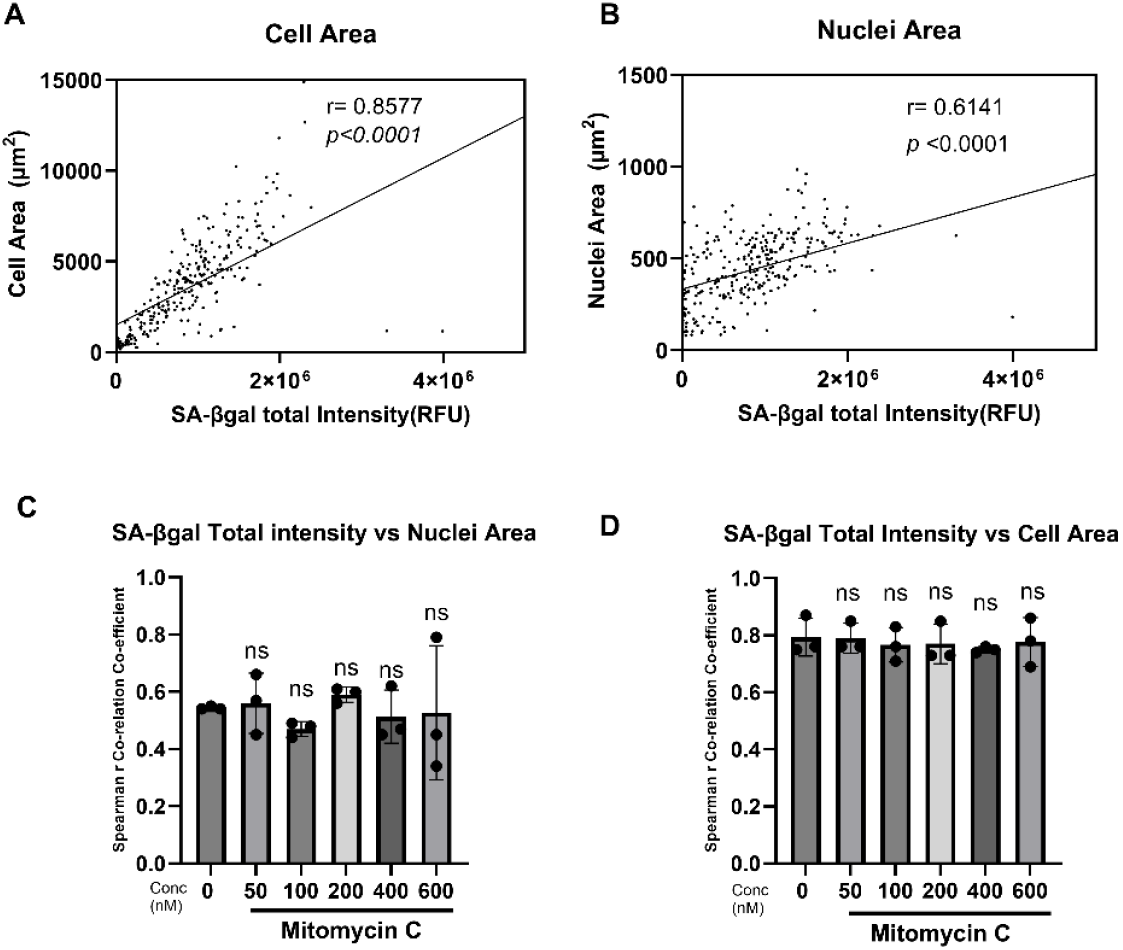
Correlation analysis of senescence-associated markers in the MMC-induced model of senescence. Representative correlation plot of SA-βgal total fluorescence intensity versus nuclear area (**A**) and ell area (**B**). Spearman correlation coefficient values for SA-βgal total fluorescence intensity versus nuclear area (**C**) and cell area (**D**). Data represent mean ± standard deviation from three independent biological replicates; ns, not significant *(p >* 0.05, ordinary one-way ANOVA with Dunnett’s post hoc test compared to the control group).

A key characteristic of senescent cells is the acquisition of a secretory phenotype, known as the senescence-associated secretory phenotype (SASP), which includes various cytokines, chemokines, and metalloproteinases. Among these, IL-6 is a major cytokine that has been extensively studied and consistently found to be elevated in senescent cells. We investigated IL-6 expression in our accelerated senescence model (Fig. 4A) and observed a three-fold increase in IL-6 expression in cells treated with 200 nM MMC (Fig. 4B). However, significant heterogeneity in IL-6 expression was evident, as indicated by the large error bars in Fig. 4B.

**Figure 4.**
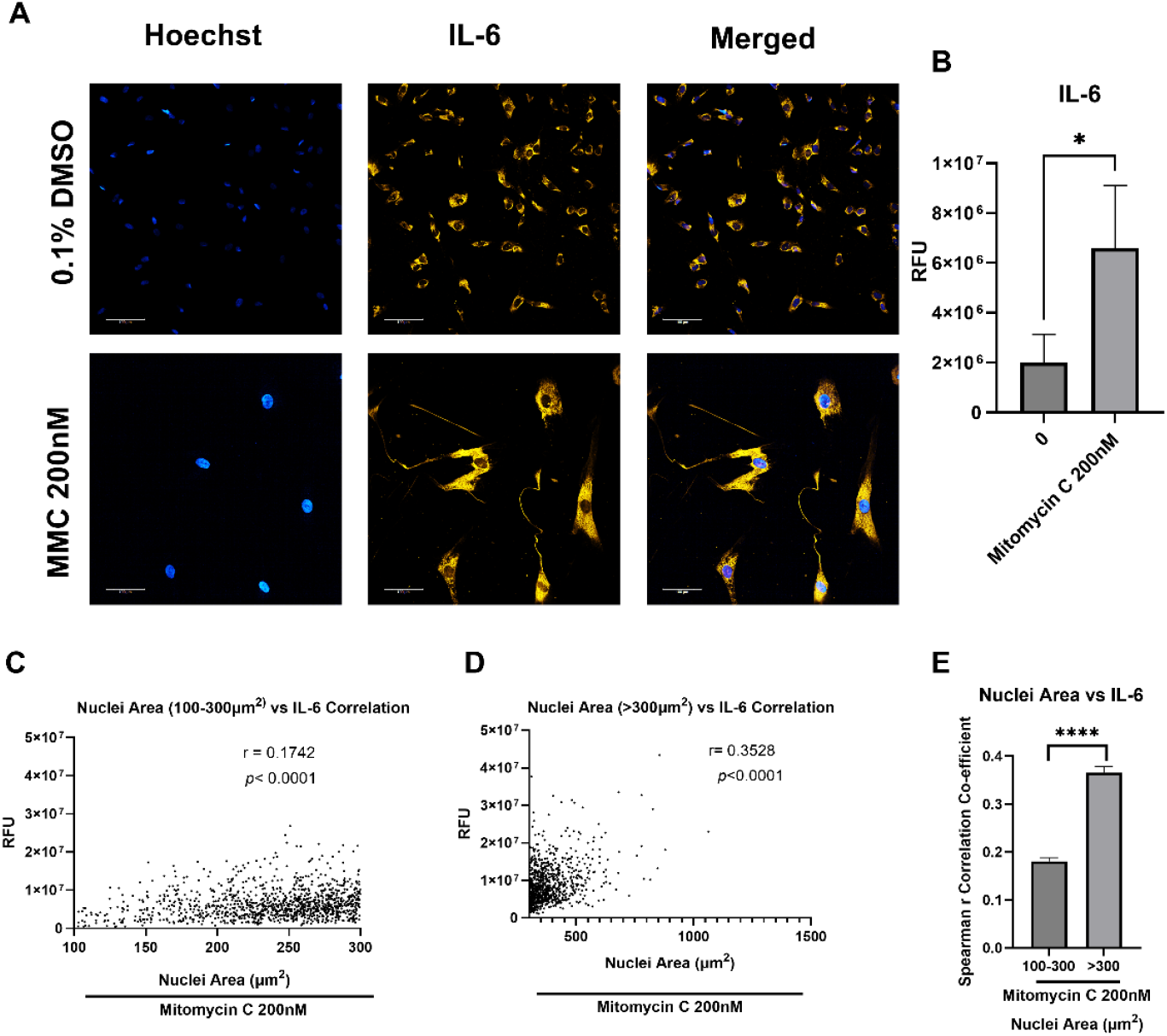
IL-6 expression in the HDF model of MMC-induced senescence. **(A)** Representative images of IL-6 expression in HDFs treated with vehicle and 200 nM MMC with nuclei labelled with Hoechst (blue) and IL-6 labelled (yellow); merged image (Scale bar: 10 µm). Images were captured using the Opera Phoenix plus™ at 20X magnification. **(B)** Average IL-6 expression in HDFs treated with vehicle or 200 nM MMC. Error bars represent the mean ± standard deviation from three independent biological replicates (Simple unpaired t-test compared to the control (0) group \**p* < 0.05). Representative correlation graph of nuclear area (100-300 µm^2^) versus IL-6 (RFU) **(C)** and nuclear area (> 300 µm^2^) versus IL-6 (RFU) **(D). (E)** Average Spearman correlation coefficient (r) of nuclear area subpopulations versus IL-6 fluorescence in HDFs treated with 200 nM MMC.

To further explore this heterogeneity, we examined the correlation between nuclear area (100-300 µm^2^) (Fig. 4C) and > 300 µm^2^ (Fig. 4D) with IL-6 expression levels. This analysis was performed based on our previous finding (Fig. 2B), which showed that the nuclear area in vehicle-treated HDFs was < 300 µm^2^. Our analysis revealed a statistically significant increase in the Spearman correlation coefficient for cells with larger nuclei (> 300 µm^2^) compared to those with nuclear areas between 100-300 µm^2^. This suggests that the expression of senescence biomarkers is associated with the pro-inflammatory secretory phenotype of the cells.

To further investigate the heterogeneity of senescence biomarkers, we examined the effects of rapamycin, a well-known senomorphic (24), on these markers. Rapamycin treatment significantly decreased the single-cell mean levels of SA-βgal fluorescence (Fig. 5A), nuclear area (Fig. 5B), and nuclear P21 fluorescence (Fig. 5C). Sub-population analysis of these biomarkers revealed a rightward shift in values with MMC treatment compared to the control, while rapamycin reduced this rightward shift (Fig. 5D-F). Applying the induction threshold method showed a statistically significant reduction in senescence biomarkers in the rapamycin-treated group (Fig. 5G-I).

**Figure 5.**
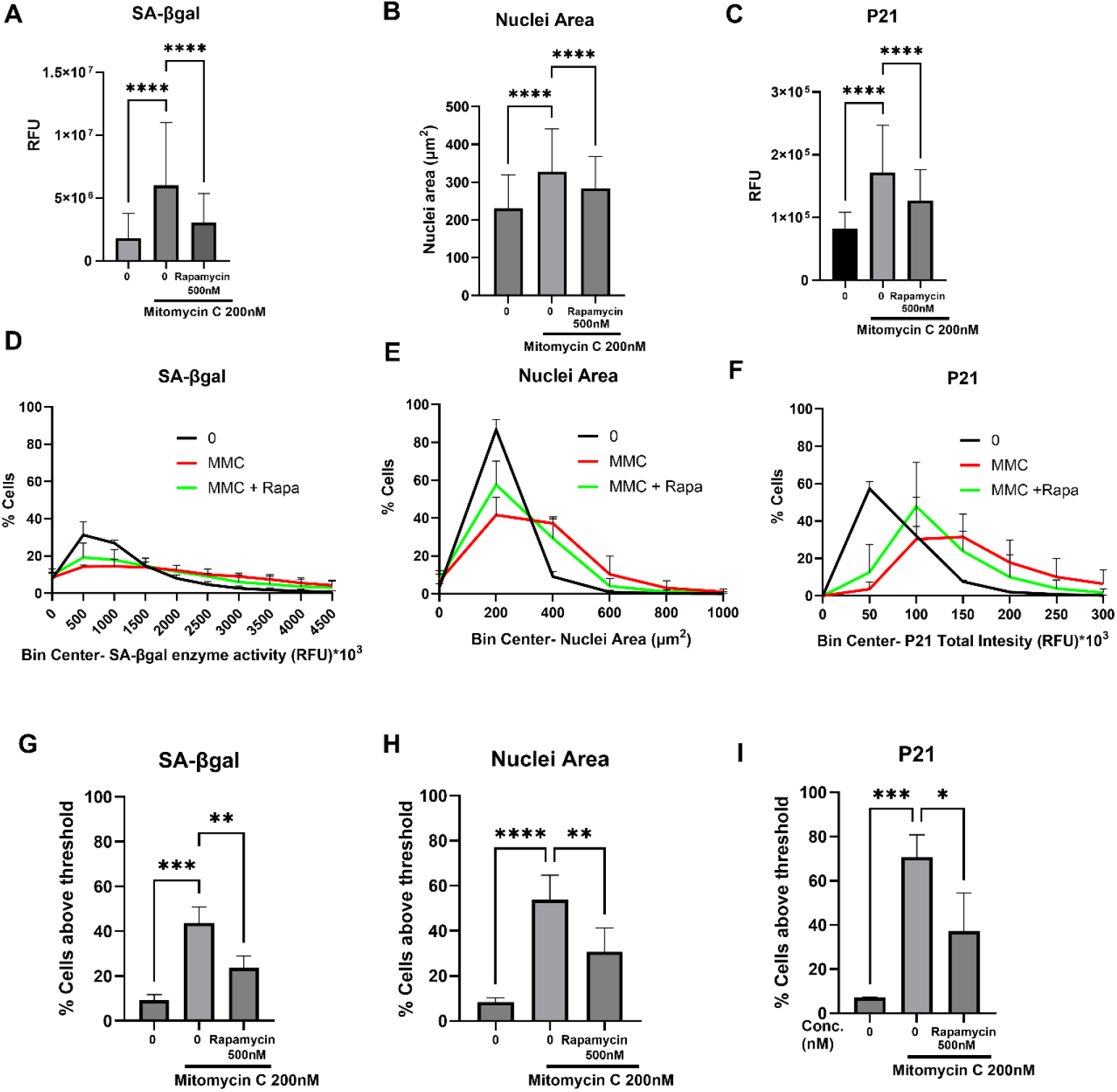
Assessment of senescence biomarkers in the MMC-induced senescence model with rapamycin treatment. Average total fluorescence intensity of SA-βgal (**A**), nuclear area (**B**) and P21 (**C**) in HDFs treated with either MMC or MMC plus rapamycin. Data represent the mean ± SD from n = 3 biological replicates; *** *p* < 0.001, **** *p* < 0.0001 (ordinary one-way ANOVA compared to the MMC 200 nM treated group). Sub-population analysis of SA-βgal (**D**), nuclear area (**E**), and P21 (**F**) in HDFs treated with either MMC or MMC plus rapamycin. Percentage of cells having increased expression of SA-βgal (**G**), nuclear area (**H**), and P21 (**I**) in HDFs treated either with MMC or MMC plus rapamycin using the induction threshold method. Data represents the mean ± SD from n = 3 biological replicates; \**p* < 0.05, \*\**p <* 0.01, *** *p* < 0.001, **** *p* < 0.0001 (ordinary one-way ANOVA compared to the MMC 200 nM treated group).

To determine if the effects of rapamycin are specific to MMC-induced senescence, we assessed senescence biomarkers in a D-galactose-induced model of senescence that we had established (Supplementary Fig. S1 and S2). We observed a trend toward reduced nuclear P21 fluorescence intensity (Fig. 6A) and a significant decrease in average nuclear and cell areas (Fig. 6B & C). Sub-population analysis revealed a rightward shift in the distribution curves of senescence biomarkers in HDFs cultured in D-galactose media compared to those cultured in glucose media (Fig. 6D-F). For nuclear P21 expression, rapamycin slightly shifted the curve leftward (Fig. 6D). Interestingly, for nuclear and cell areas, rapamycin-treated HDFs in D-galactose media showed a shift in the curve back to control levels observed in HDFs cultured in glucose media (Fig. 6E & F). Using the induction threshold method, we observed a two-fold reduction in nuclear P21 expression in rapamycin-treated cells compared to D-galactose-only treated cells (Fig. 6G), while nuclear and cell areas exhibited a three-fold reduction relative to the D-galactose group (Fig. 6H & I).

**Figure 6.**
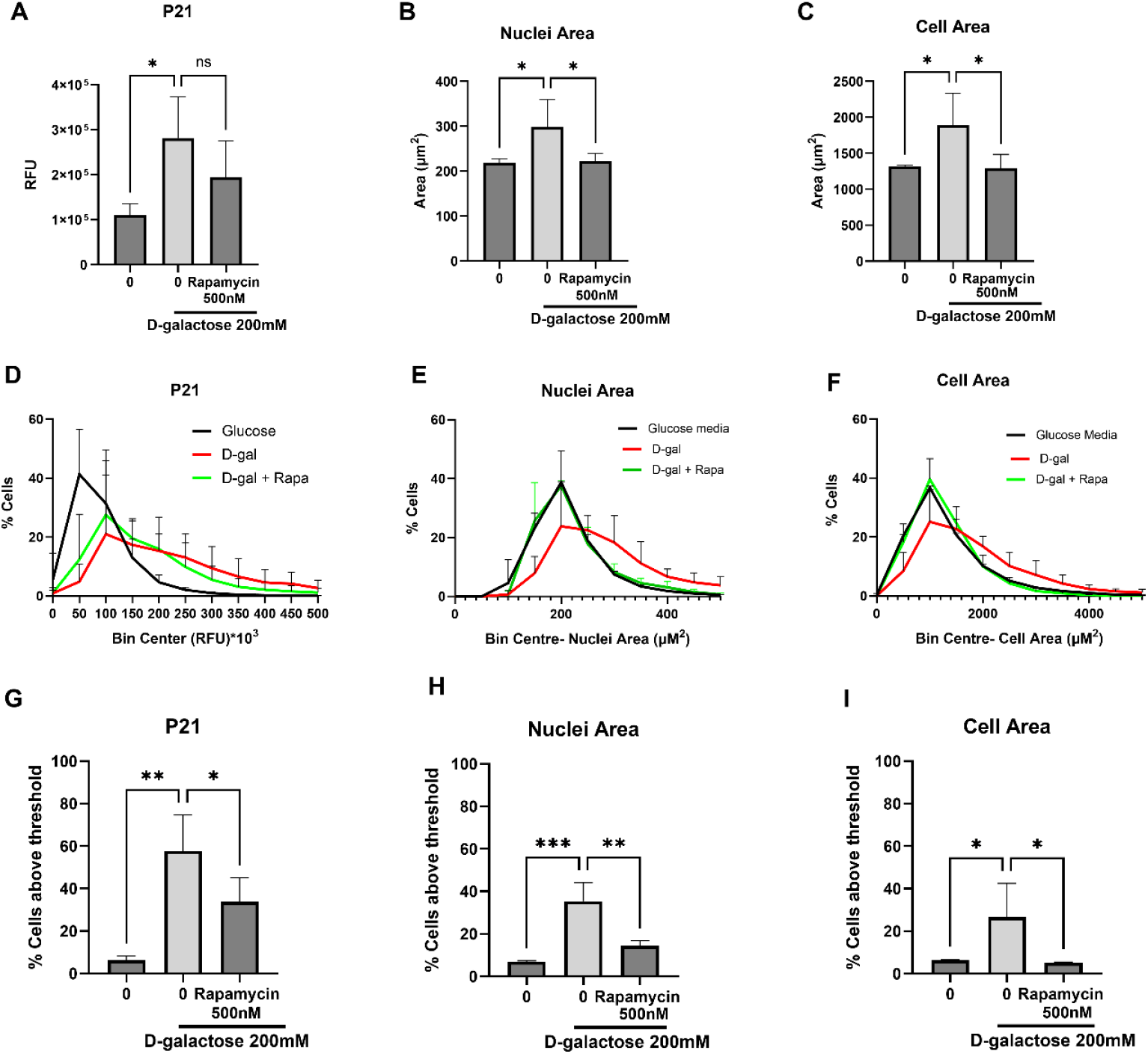
Assessment of senescence biomarkers in D-galactose-induced model of senescence with rapamycin treatment. Average total fluorescence intensity of P21 (**A**), nuclear area (**B**), and cell area (**C**) in HDFs cultured in either glucose media, 200 mM D-galactose media, or 500 nM rapamycin plus 200 mM D-galactose media. Data represents the mean ± SD from n = 3 biological replicates; * *p* < 0.05, (ordinary one-way ANOVA compared to the 0 (D-galactose media) group). Sub-population analysis of P21 (**D**), nuclear area (**E**), and cell area (F) in HDFs cultured in either glucose media, 200 mM D-galactose media, or 500 nM rapamycin plus 200 mM D-galactose media. Percentage of cells with increased expression of P21 (**G**), nuclear area (**H**), and cell area (**I**), in HDFs cultured in either glucose media, 200 mM D-galactose media, or 500 nM rapamycin plus 200 mM D-galactose media, using the induction threshold method. Data represents the mean ± SD from n = 3 biological replicates; \**p* < 0.05, \*\**p <* 0.01, *** *p* < 0.001, (ordinary one-way ANOVA compared to the 0 (D-galactose media) group).

## Discussion

This study explored the heterogeneous expression of senescence biomarkers using single-cell fluorescence microscopy, introducing a novel approach to analyse and quantify these biomarkers to account for diverse cell populations within a primary cell culture. At present, no single, universally accepted marker of senescence is available. Traditionally, SA-βgal is assessed in fixed cells through histochemical analysis, which does not allow for quantitative studies of expression levels in senescent cells (17). Fibroblasts, one of the gold-standard models for studying cellular senescence, are not homogeneous in their replicative state, even at early passages. With each passage, a subset of cells progresses towards cellular senescence, leading to a heterogeneous population at different stages of the senescence programme. This heterogeneity presents challenges in robustly assessing the efficacy of senotherapeutics for drug screening.

In this study, live-cell imaging was used to reliably assess three senescence biomarkers including increased SA-βgal activity and two morphological features, nuclear area and cell area, in live cells. SA-βgal activity and nuclear area, in addition to increasing with D-galactose and MMC treatment, displayed notable heterogeneity at the single-cell level, revealing distinct sub-populations within treated cells. Our findings indicate that in both chemotherapy-induced and D-galactose models of senescence, rapamycin treatment selectively impacted specific sub-populations of senescence biomarkers. This highlights the importance of assessing heterogeneity within these biomarkers to better understand senescence dynamics. The physiological significance of these sub-populations warrants further investigation.

The current data indicate a strong correlation between nuclear area and cell area with SA-βgal activity. This suggests that a simple assessment of nuclear or cell area can be used interchangeably to determine senescence induction. One major disadvantage of SA-βgal is its overexpression in confluent cells, which can lead to false-positive results. This drawback can be addressed by assessing nuclear size, which is not affected by the confluency of fibroblasts.

Our data also reveal heterogeneity in the increase in nuclear area. We hypothesise that this may be due to a correlation between nuclear size and the SASP level, as evidenced by the stronger correlation of IL-6 fluorescence levels with increasing nuclear area. However, further studies are required to confirm this association.

Our data show varying levels of nuclear P21 expression in senescent cells. Recent studies have revealed that P21 is involved in cell fate decisions and induces an independent biosecretome(24). Thus, these observed differences in P21 levels may suggest that, depending on P21 expression levels, cells may have distinct fates or an altered regulation of SASP secretion. Interestingly, a small population of MMC-treated fibroblasts incorporated EdU into their nuclei, although these cells did not replicate over time. We hypothesise that this may be due to either paracrine effects from neighbouring cells or these cells’ limited ability to repair and synthesise DNA, though not enough to proliferate into daughter cell populations.

In addition, P21 and TP53 expression is involved in mediating apoptosis (25). Higher levels of P21 render cells more resistant to apoptosis, as senescent cells are known to exhibit anti-apoptotic properties (25). Therefore, drugs that reduce P21 levels might sensitise cells to senolytic therapies targeting the p53-P21 axis in senescence.

The induction threshold method described in this paper demonstrates that, due to varying expression levels of senescence biomarkers, analysing the entire population as a whole and averaging the data may not be the most effective approach for assessing senescence biomarkers. Indeed, we show that western blotting revealed approximately a two-fold increase in P21 expression, similar to the average values observed using the fluorescence microscopy-based method. However, the induction threshold method indicated a robust six-to seven-fold change, suggesting that averaging data might mask the efficacy of senotherapeutic compounds. Thus, this new method provides a more robust way to identify novel senotherapeutic drugs.

In addition to these advantages, the present study demonstrated that the induction threshold method can be applied to different senescence inducers, acting through various mechanisms, such as D-galactose-induced senescence. This is a key point of differentiation, as many senotherapeutic effects are limited to specific senescence effectors and cell types.

The present study has demonstrated that the induction threshold method can be applied to senotherapeutic drug discovery, as evidenced by the senotherapeutic effect of rapamycin, a drug that is currently in phase 3 clinical trials for longevity extension and known as a senomorphic (24). We show that rapamycin significantly decreased all the aforementioned biomarkers of senescence. The main advantage of this method is that, by analysing biomarkers at the single-cell level, the pharmacological effect size of the drug can be robustly demonstrated *in vitro*. Thus, this single-cell analysis of senescence biomarkers is suitable for medium-to high-throughput drug discovery screens.

In conclusion, this study highlights the heterogeneous expression of senescence biomarkers within a cell population using fluorescence microscopy and presents an alternative single-cell method for the robust discovery of senotherapeutic drug candidates.

## ACKNOWLEDGEMENTS

V.S. was supported by a Tasmanian Graduate Research Scholarship. The experimental component of this research was funded by the University of Tasmania and Monash University. The authors acknowledge the facilities, and scientific and technical assistance of the Monash Biomedicine Discovery Institute (BDI) Organoid Program and Monash Micro Imaging at Monash University, Victoria, Australia.

## AUTHORS’ CONTRIBUTIONS

V.S. and I.A. conceived the study and designed the experiments. V.S. performed all the experiments except the western blot which was performed by J.T. V.S, C.C. and I.A wrote the manuscript. I.A. and N.G. supervised the project. I.A, N.G., K.B., and Y.W. reviewed the manuscript and provided intellectual input.

## References

1. Hayflick L, Moorhead PS. The serial cultivation of human diploid cell strains. Experimental Cell Research. 1961;25(3):585–621.

2. Yousefzadeh MJ, Flores RR, Zhu Y, Schmiechen ZC, Brooks RW, Trussoni CE, et al. An aged immune system drives senescence and ageing of solid organs. Nature. 2021;594(7861):100–5.

3. Shay JW, Wright WE. Hayflick, his limit, and cellular ageing. Nature reviews Molecular cell biology. 2000;1(1):72–6.

4. Herranz N, Gil J. Mechanisms and functions of cellular senescence. The Journal of clinical investigation. 2018;128(4):1238–46.

5. Childs BG, Baker DJ, Kirkland JL, Campisi J, Van Deursen JM. Senescence and apoptosis: dueling or complementary cell fates? EMBO reports. 2014;15(11):1139–53.

6. Lee S, Lee J-S. Cellular senescence: A promising strategy for cancer therapy. BMB reports. 2019;52(1):35.

7. Gorgoulis V, Adams PD, Alimonti A, Bennett DC, Bischof O, Bishop C, et al. Cellular senescence: defining a path forward. Cell. 2019;179(4):813–27.

8. Domen A, Deben C, Verswyvel J, Flieswasser T, Prenen H, Peeters M, et al. Cellular senescence in cancer: clinical detection and prognostic implications. Journal of Experimental & Clinical Cancer Research. 2022;41(1):360.

9. McHugh D, Gil J. Senescence and aging: Causes, consequences, and therapeutic avenues. J Cell Biol. 2018;217(1):65–77.

10. Di Micco R, Krizhanovsky V, Baker D, d’Adda di Fagagna F. Cellular senescence in ageing: from mechanisms to therapeutic opportunities. Nature Reviews Molecular Cell Biology. 2021;22(2):75–95.

11. Coppé JP, Desprez PY, Krtolica A, Campisi J. The senescence-associated secretory phenotype: the dark side of tumor suppression. Annu Rev Pathol. 2010;5:99–118.

12. Davalos AR, Coppe J-P, Campisi J, Desprez P-Y. Senescent cells as a source of inflammatory factors for tumor progression. Cancer and Metastasis Reviews. 2010;29:273–83.

13. Amaya-Montoya M, Pérez-Londoño A, Guatibonza-García V, Vargas-Villanueva A, Mendivil CO. Cellular senescence as a therapeutic target for age-related diseases: a review. Advances in therapy. 2020;37:1407–24.

14. Schwartz GK, Shah MA. Targeting the cell cycle: a new approach to cancer therapy. J Clin Oncol. 2005;23(36):9408–21.

15. He S, Sharpless NE. Senescence in health and disease. Cell. 2017;169(6):1000–11.

16. HAmaral JD, Xavier JM, Steer CJ, Rodrigues CM. The role of p53 in apoptosis. Discovery medicine. 2010;9(45):145–52.

17. Dimri GP, Lee X, Basile G, Acosta M, Scott G, Roskelley C, et al. A biomarker that identifies senescent human cells in culture and in aging skin in vivo. Proc Natl Acad Sci U S A. 1995;92(20):9363–7.

18. de Mera-Rodríguez JA, Álvarez-Hernán G, Gañán Y, Martín-Partido G, Rodríguez-León J, Francisco-Morcillo J. Is senescence-associated β-galactosidase a reliable in vivo marker of cellular senescence during embryonic development? Frontiers in cell and developmental biology. 2021;9:623175.

19. Yang N-C, Hu M-L. The limitations and validities of senescence associated-β-galactosidase activity as an aging marker for human foreskin fibroblast Hs68 cells. Experimental Gerontology. 2005;40(10):813–9.

20. Rufini A, Tucci P, Celardo I, Melino G. Senescence and aging: the critical roles of p53. Oncogene. 2013;32(43):5129–43.

21. Azman KF, Zakaria R. D-Galactose-induced accelerated aging model: an overview. Biogerontology. 2019;20(6):763–82.

22. Wang R, Yu Z, Sunchu B, Shoaf J, Dang I, Zhao S, et al. Rapamycin inhibits the secretory phenotype of senescent cells by a Nrf2-independent mechanism. Aging Cell. 2017;16(3):564–74.

23. Severino J, Allen RG, Balin S, Balin A, Cristofalo VJ. Is β-Galactosidase Staining a Marker of Senescence in Vitro and in Vivo? Experimental Cell Research. 2000;257(1):162–71.

24. Sturmlechner I, Zhang C, Sine CC, van Deursen EJ, Jeganathan KB, Hamada N, et al. P21 produces a bioactive secretome that places stressed cells under immunosurveillance. Science. 2021;374(6567):eabb3420.

25. Yosef R, Pilpel N, Papismadov N, Gal H, Ovadya Y, Vadai E, et al. P21 maintains senescent cell viability under persistent DNA damage response by restraining JNK and caspase signaling. Embo j. 2017;36(15):2280–95.

