## Supplementary Figures for "Single-cell fluorescence imaging reveals heterogeneity in senescence biomarkers and identifies rapamycin-responsive sub-populations"

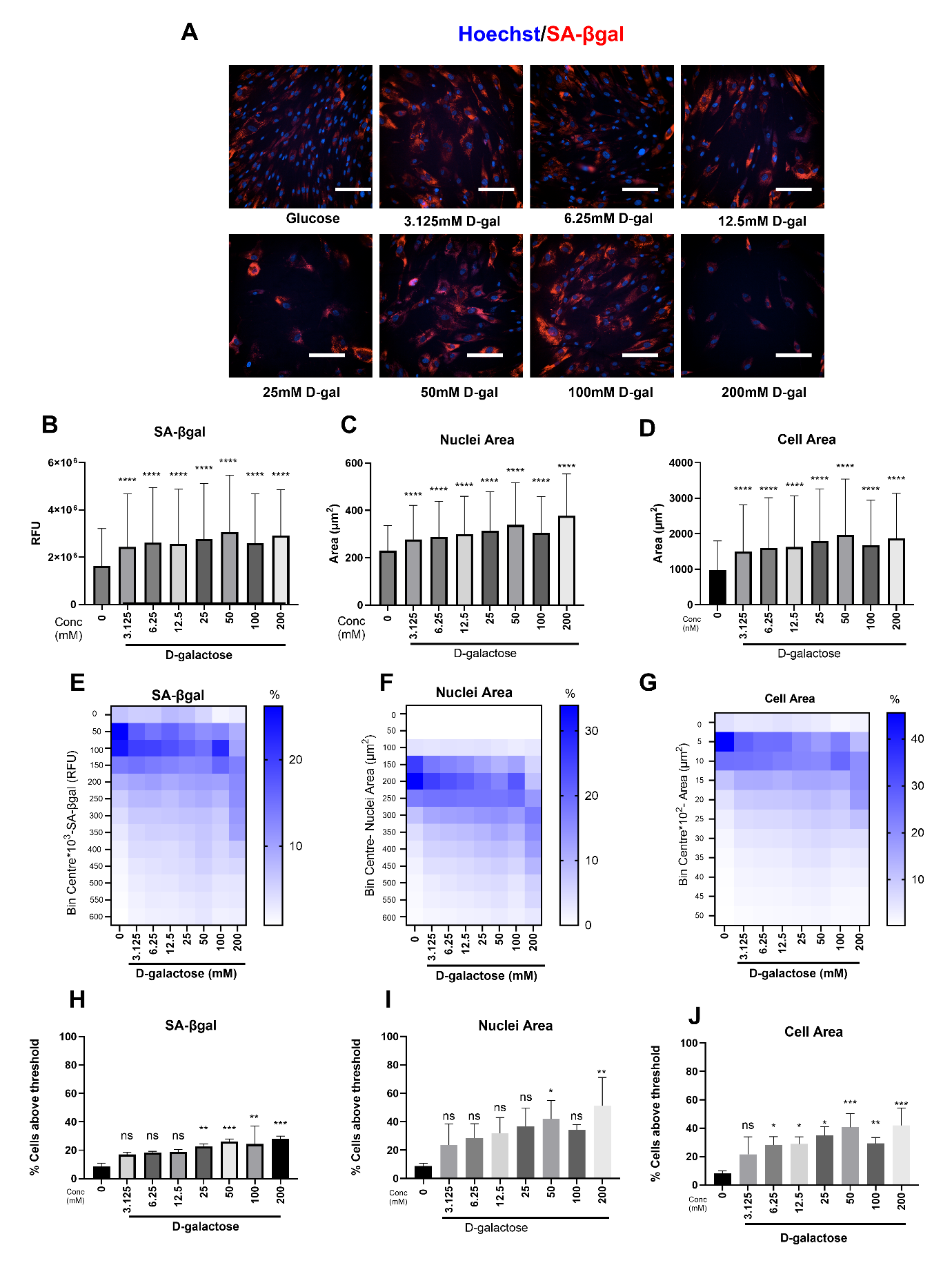

**Figure S1. Senescence markers in D-galactose (D-gal)-induced senescence.** (**A**) Representative image of HDFs cultured in glucose-containing media and various concentrations of D-galactose (3.125–200 mM) with nuclei stained with Hoechst (blue) and SA-βgal (red); scale bar: 200 µm. Images were taken using the Opera Phenix™ Plus at 20x magnification. Average total fluorescence intensities of SA-βgal (**B**), average nuclear area (**C**), and cell area (**D**) in HDFs cultured in D-galactose media; error bars represent mean ± standard deviation from one experiment with three replicates. *****p* < 0.0001 (one-way ANOVA with Kruskal-Wallis test compared to the control group). Individual cell-derived heatmaps of SA-βgal (**E**), nuclear area (**F**), and cell area (**G**) of HDFs cultured in D-galactose media. Percentage of cells with SA-βgal fluorescence total intensity (**H**), nuclear area (**I**), and cell area (**J**) above the threshold set for cells cultured in glucose-containing media; ns, not significant (*p* > 0.05), **p* < 0.05, ***p* < 0.01, ****p* < 0.001 (ordinary one-way ANOVA compared to control group).

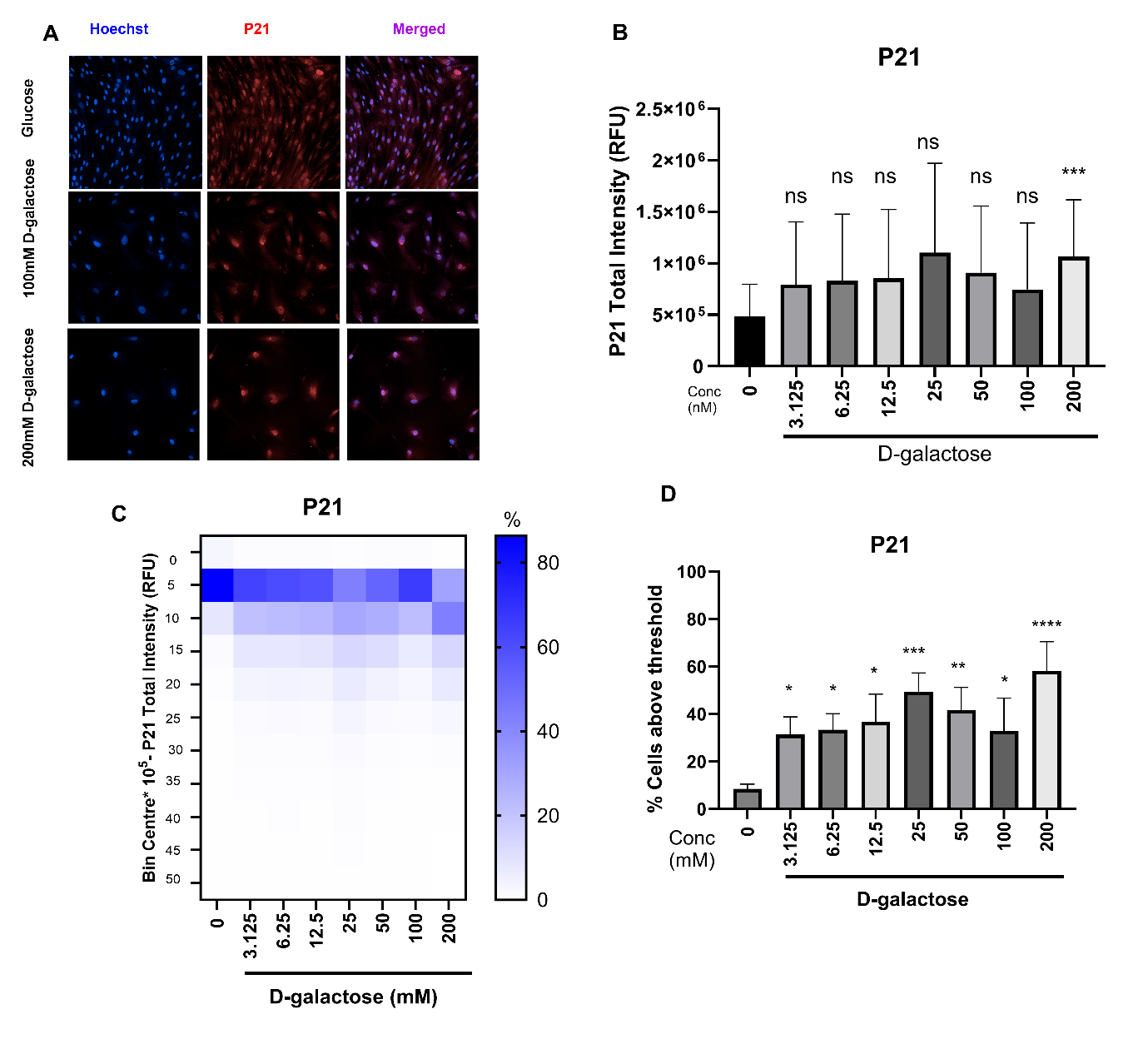

**Figure S2. Expression of P21 in D-galactose-induced senescence.** (**A**) Representative images of P21 in HDFs cultured in D-galactose-containing media with nuclei stained with Hoechst (blue), P21 (red), and merged; scale bar: 200 µm. Images were taken using the Opera Phenix™ Plus at 20x magnification. (**B**) Total fluorescence intensity of nuclear P21 in HDFs cultured in D-galactose-containing media; *****p* < 0.0001 (one-way ANOVA with Kruskal-Wallis test compared to the control group). (**C**) Single-cell histogram of total fluorescence intensity of nuclear P21 in HDFs cultured in D-galactose-containing media. (**D**) Percentage of cells positive for P21 expression in HDFs cultured in D-galactose-containing media, assessed using the induction threshold method; **p* < 0.05, ***p* < 0.01, ****p* < 0.001, *****p* < 0.0001 (ordinary one-way ANOVA compared to control group).
